# Lithium ameliorates Cornelia de Lange syndrome associated phenotypes in experimental models

**DOI:** 10.1101/2020.07.15.204628

**Authors:** Paolo Grazioli, Chiara Parodi, Milena Mariani, Daniele Bottai, Elisabetta Di Fede, Aida Zulueta, Laura Avagliano, Anna Cereda, Romano Tenconi, Jolanta Wierzba, Raffaella Adami, Maria Iascone, Paola Francesca Ajmone, Thomas Vaccari, Cristina Gervasini, Angelo Selicorni, Valentina Massa

**Affiliations:** Department of Health Sciences, Università degli Studi di Milano, Milan, Italy; UOC Pediatria, ASST Lariana, Como, Italy; “Aldo Ravelli” Center for Neurotechnology and Experimental Brain Therapeutics, Università degli Studi di Milano, Milan, Italy; Department of Pediatrics - ASST Papa Giovanni XXIII, Bergamo, Italy; Department of Pediatrics, University of Padova, Padova, Italy; Department of Pediatrics and Internal Medicine Nursing, Department of Rare Disorders, Medical University of Gdansk Poland; Child and Adolescent Neuropsychiatric Unit, Fondazione IRCCS Cà Granda Ospedale Maggiore Policlinico, Milan, Italy; Department of Biosciences, Università degli Studi di Milano, Milano, Italy

**Author notes:** These authors contributed equally to this work (Paolo Grazioli, Chiara Parodi, Milena Mariani). **Correspondence** and requests for materials should be addressed to V.M.

## Abstract

Cornelia de Lange Syndrome (CdLS) is a rare developmental disorder affecting a multitude of organs including the central nervous system, inducing a variable neurodevelopmental delay. CdLS malformations derive from deregulation of developmental pathways, inclusive of the canonical WNT pathway.

We have evaluated MRI anomalies and behavioral and neurological clinical manifestations in CdLS patients. Importantly, we observed in our cohort a significant association between behavioral disturbance and structural abnormalities in brain structures of hindbrain embryonic origin.

Considering the cumulative evidence on the cohesin-WNT-hindbrain shaping cascade, we have explored possible ameliorative effects of chemical activation of the canonical WNT pathway with lithium chloride in different models: (I) *Drosophila melanogaster* CdLS model showing a significant rescue of mushroom bodies morphology in the adult flies; (II) mouse neural stem cells restoring physiological levels in proliferation rate and differentiation capabilities toward the neuronal lineage; (III) lymphoblastoid cell lines from CdLS patients and healthy donors restoring cellular proliferation rate and inducing the expression of *CyclinD1*. This work supports a role for WNT-pathway regulation of CdLS brain and behavioral abnormalities and a consistent phenotype rescue by lithium in experimental models.

## INTRODUCTION

Cornelia De Lange Syndrome (CdLS, OMIM #122470, #300590, #610759, #614701, #300882) is a rare genetic disorder affecting a variety of organs, including the Central Nervous System (CNS). This syndrome is mainly caused by dominant autosomal or X-linked *de novo* mutations and it is a genetically and clinically heterogeneous disorder.

The recent International Consensus Statement (1) has defined the phenotypes classified as CdLS as a spectrum including the classic CdLS phenotype with or without a pathogenic variant in a gene involved in cohesin functioning, as well as individuals with a non-classic CdLS phenotype with a pathogenic variant in a cohesin function-relevant gene. The spectrum does not include those patients with a variant in a cohesin gene without the CdLS phenotype.

The prevalence of CdLS is estimated to be 1:10.000-30.000 newborns but, most likely this represents an underestimation, as milder cases may not be recognized (2).

The features of this disorder vary widely among affected individuals and range from relatively mild to severe. CdLS is characterized by slow growth, abnormalities of the bones in the arms, hands and fingers, and disorders in the gastro-intestinal tract. Patients present distinctive facial features, including arched eyebrows, long eyelashes, low-set ears, small and widely spaced teeth, and a small and upturned nose. CdLS patients are characterized by intellectual disabilities and behavior of the autism-spectrum indicating neural development alterations (3, 4).

CdLS is primarily caused by mutations in one of 5 genes: *NIPBL, SMC1A, SMC3, RAD21*, and *HDAC8* (1, 4). These genes encode for proteins of the cohesin complex that is a multimeric system, highly conserved in the course of cellular evolution from the most primitive life forms to human cells (5–7). Cohesins are essential Structural Maintenance of Chromosomes (SMC) protein-containing complexes that interact with chromatin and modulate chromatin organization. Cohesins mediate sister chromatid cohesion and cellular long-distance chromatin interactions affecting genome maintenance and gene expression (8–10).

The current hypothesis regarding CdLS pathogenesis is that malformations arise from a deregulation of developmental pathways (1) and in this context, ours and other previous studies (4, 11–13) have shown that the canonical WNT pathway is perturbed.

The WNT pathway regulates cell-cell signaling by means of two branches. A canonical pathway acts through frizzled family receptors, which relay the signals to the intracellular transducer protein Dishevelled and regulates the expression of key developmental target genes (14). In vertebrates, a non-canonical WNT pathway (also called β-catenin-independent pathway) is known to regulate both cell polarity and dorsal mesodermal cell movements during convergent extension, and later during neural tube closure (15, 16).

WNT signaling is involved in numerous events in animal development and in the maintenance of adult tissue homeostasis by regulating cell proliferation, differentiation (including the proliferation of stem cells and the specification of the neural crest), migration, genetic stability and apoptosis, as well as by maintaining adult stem cells in a pluripotent state (17).

The WNT pathway plays a fundamental role in all steps of development of the CNS. In fact, WNT signaling alterations have been associated with a plethora of CNS abnormalities (4). We have previously shown that the canonical WNT pathway is perturbed in CdLS models and underlies the observed cellular and developmental alterations. It is well known that lithium, a clinically effective therapeutic agent for bipolar disorders (18), also recently used for amyotrophic lateral sclerosis (19), is a specific inhibitor of the conserved WNT transducer GSK-3β (20, 21). Indeed, our group recently reported that lithium activates the WNT pathway and is able to rescue phenotypes of *nipbl* and *smc1a* knock-down zebrafish mutants. Moreover, we observed that lithium-dependent activation of WNT pathway was able to rescue phenotypes of *nipbl* and *smc1a* knock-down zebrafish mutants (11, 12).

Based on these data, our working model for CdLS is that the canonical WNT pathway is downregulated as a consequence of a haploinsufficiency of cohesin genes. Such downregulation leads to lower expression of *CylinD1* which, in turn, leads to an increase in cell death at specific times and in specific tissues of the developing embryo (8, 11, 12).

Our experimental data was placed in context with CdLS patient clinical findings: in particular, with radiological anomalies and behavioral and neurological phenotypes. Thus, herein we decided to investigate the possible ameliorative effects of treatment with lithium, exploiting *in vivo* and *in vitro* CdLS models.

## RESULTS

### Cognitive and behavioral deficits in CdLS patients are associated with abnormalities of hindbrain-derived structures

The relationship between WNT pathway and cohesin complex genes haploinsufficiency has been previously demonstrated especially in the case of hindbrain embryonic development (4). For this reason, we sought to ascertain if alterations in cognitive, neurological and behavioral aspects of CdLS patients could be associated with morphological abnormalities in hindbrain-derived structures (22). To this end, we evaluated Magnetic Resonance Imaging (MRI) images of a CdLS cohort (Supp. Fig. 1B). Among the 66 brain MRI, 31 showed brain abnormalities, half of which exhibited more than one anomaly (Fig. 1A; Supp. Fig. 2A). The cerebellum and cisterna magna were the most frequently affected structures among recorded abnormalities (both affecting 25% of patients) and the most frequently coexisting anomalies. Subdividing MRI data according to individual behavioral assessment, we observed a significant correlation between CNS abnormalities and autism spectrum disorder (ASD) (Fig. 1B-C). Subdividing MRI data according to cognitive level, we observed a significant correlation between CNS malformations and severe cognitive impairment (Supp. Fig. 2B-C). MRI findings were not associated with the presence/absence of seizure (observed in 13/66 individuals). Importantly, behaviors associated with ASD significantly correlated to abnormalities of structures derived from the hindbrain (Fig. 1C).

**Figure 1.**
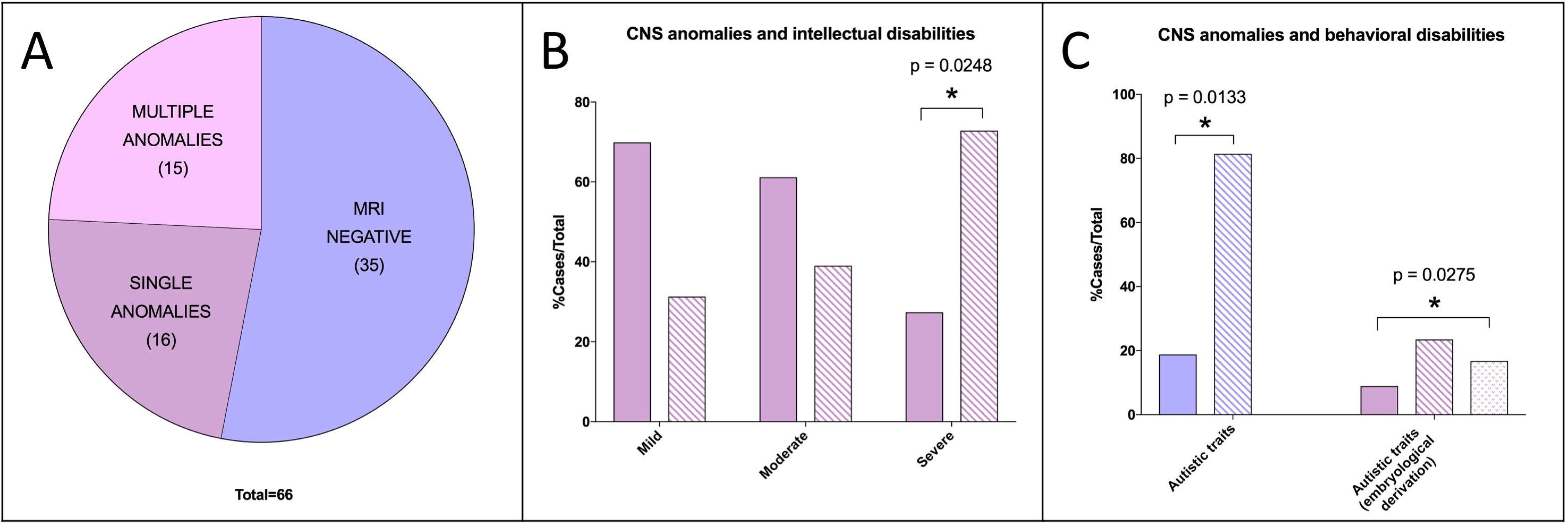
CdLS brain abnormalities in hindbrain derived structures correlates with cognitive and behavioral alterations. (A) MRI data distribution in the cohort of CdLS patients. Purple: MRI negative (35/66); mauve: MRI positive – single anomalies (16/66); pink: MRI positive – multiple anomalies (15/66). (B) CNS anomalies and intellectual disabilities. Solid mauve: MRI negative; striped mauve: MRI anomalies. (C) CNS anomalies and behavioral disabilities – autistic traits. Solid purple: MRI negative; striped purple: MRI anomalies; solid mauve: MRI negative; striped mauve: Rhombencephalon derivation; dotted mauve: Prosencephalon derivation. * p<0.05; * vs MRI negative.

### WNT activation restores correct mushroom body development in CdLS flies

To study the effects of lithium on organ development, we next moved on to an *in vivo* CdLS model. To this end, we decided to evaluate the morphology of mushroom bodies of *Drosophila melanogaster*, a well-studied specialized CNS structure involved in olfactory learning and memory in adults (Fig. 2A). Indeed, flies heterozygous for an inactivating mutation in the fly *NIPBL* ortholog gene *Nipped-B* (*Nipped-B^407^* haploinsufficient animals from here on), often display mushroom bodies anomalies such as aberrant or missing lobes (Fig. 2C), when compared to control animals (Fig. 2B and (23)). Importantly, upon rearing *Nipped-B^407^* haploinsufficient animals on food supplemented with lithium (100 mM), a statistically significant (p=0.0036) ratio of adults did not show signs of altered mushroom body development (Fig. 2D). In particular, the percentage of *Nipped-B^407^* haploinsufficient animals with abnormal mushroom body morphology decreased from 88% in the untreated sample to 30% in the treated animals (Fig. 2E). To assess whether such significant anatomic rescue was WNT-dependent, we analyzed gene expression in our experimental groups. We observed a significant increase in expression of *engrailed* (*en* Fig. 2H-I) - a known fly WNT signaling target (24), in the *Nipped-B^407^* haploinsufficient flies treated with lithium, when compared to flies reared on unsupplemented food. As expected (25), we did not observe gene expression changes in *armadillo* (*arm*, the fly ortholog of vertebrate *β-catenin*. Fig. 2F-G) both in *yw*, control strain, and in *Nipped-B^407^* haploinsufficient animals.

**Figure 2.**
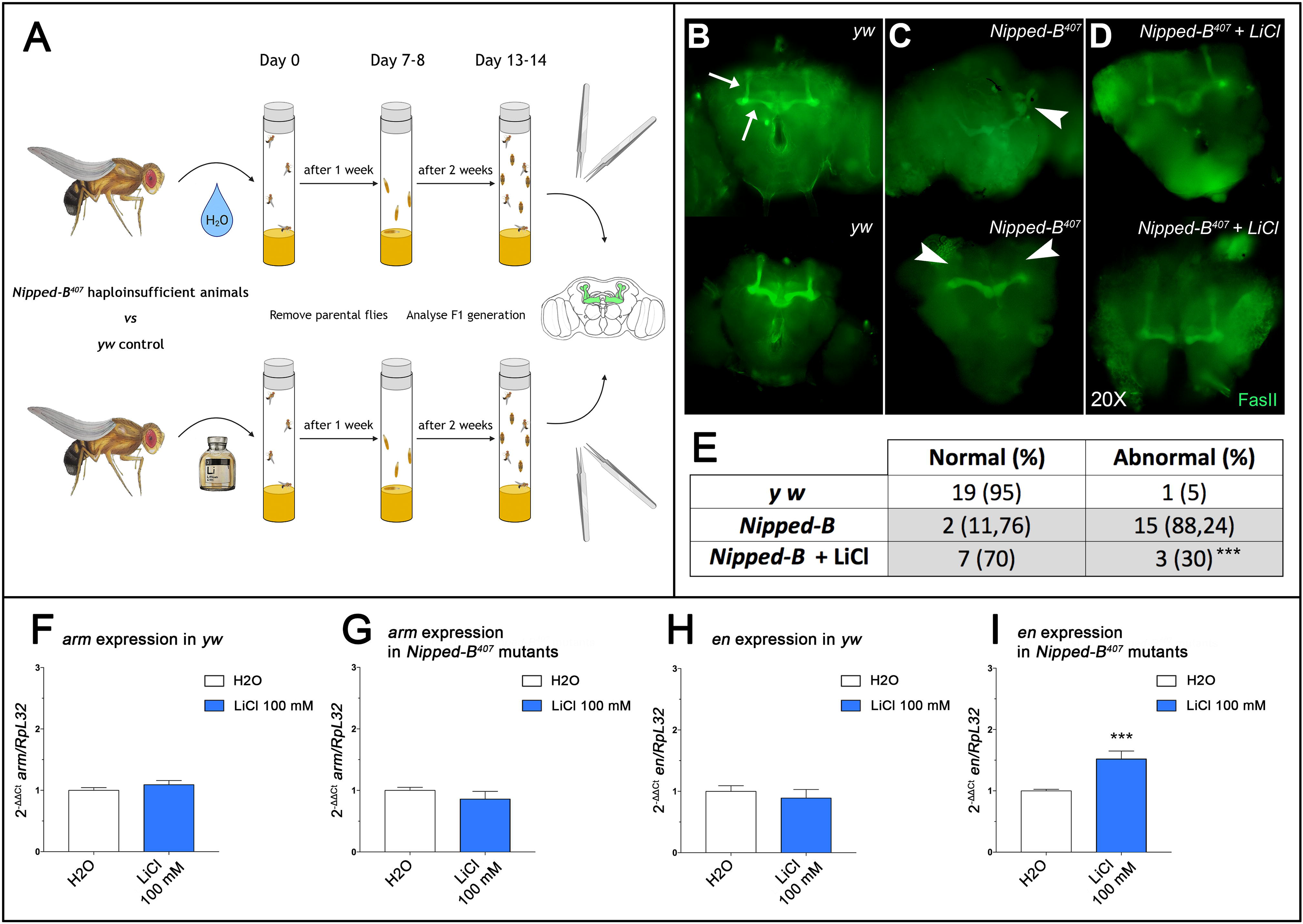
Lithium rescues mushroom bodies morphology in CdLS *D. melanogaster* through WNT activation. (A) Schematic representation of the drug-treatment protocol. (B-D) Mushroom bodies of adult animals labelled with anti-FasII are shown in green. (B) Normal morphology of mushroom bodies was observed in *yw* controls where α, β and γ lobes can be distinguished by their distinct projection patterns (arrows). (C) Abnormal mushroom bodies morphology was observed in *Nipped-B^407^* haploinsufficient adults with a twisted structure (arrowhead in the upper brain) or lacking both α lobes (arrowhead in the lower brain) as example. (D) Rescue of mushroom bodies morphology is shown in *Nipped-B^407^* haploinsufficient adults upon treatment with LiCl. (E) The table reports the total number and percentage of normal or abnormal adult mushroom bodies. Data were analyzed using Fisher’s exact test (*** p<0.005). (F-I) Histograms show *arm* (F-G) and *en* (H-I) gene expression levels in whole flies as 2^−ΔΔCt^ ± SD. Analyses of *yw* controls are shown in the panels (F-H) and analyses of the *Nipped-B^407^* mutants are shown in the panels (H-I). White: water; dark blue: LiCl 100 mM treatment. *** p<0.005; * LiCl 100 mM vs H2O.

### WNT activation restores physiological proliferation and increases neuronal differentiation in CdLS mouse neural stem cells

To study possible beneficial effects of lithium treatments in CdLS models, we exploited mouse neural stem cells (NSCs) treated with a selective inhibitor of HDAC8 enzymatic activity (PCI34051), to mimic the molecular defects observed in patients (1, 7, 26). We have previously shown that PCI34051 was able to alter the rate of mouse NSCs proliferation and differentiation (13). Thus, we tested the effects of lithium to prevent the detrimental effects of HDAC8 inhibition (Fig. 3A, 3C). We confirmed that PCI34051 exposure induces a drastic reduction of NSCs proliferation. However, simultaneous exposure to 3 mM lithium reduced the adverse effects of HDAC8 inhibition, inducing a significant increase in NSC proliferation (Fig. 3A, 3C). The protective effect of lithium was also tested using *Nipbl* knockdown by siRNA. Such knockdown caused a significant reduction of NSCs proliferation as expected. However, lithium exposure was able to completely correct the effect of *Nipbl* knockdown, leading to a significantly increased proliferation when compared to siRNA treatment, ultimately restoring proliferation levels to those observed in control NSCs (AllStar/control siRNA or AllStar/control siRNA + lithium) (Fig. 3B, 3D). In order to confirm that the ameliorative effects of lithium were mediated by WNT activation, we analyzed the expression of the Wnt target *Ccnd1* and observed a significant increase (Fig. 3E).

**Figure 3.**
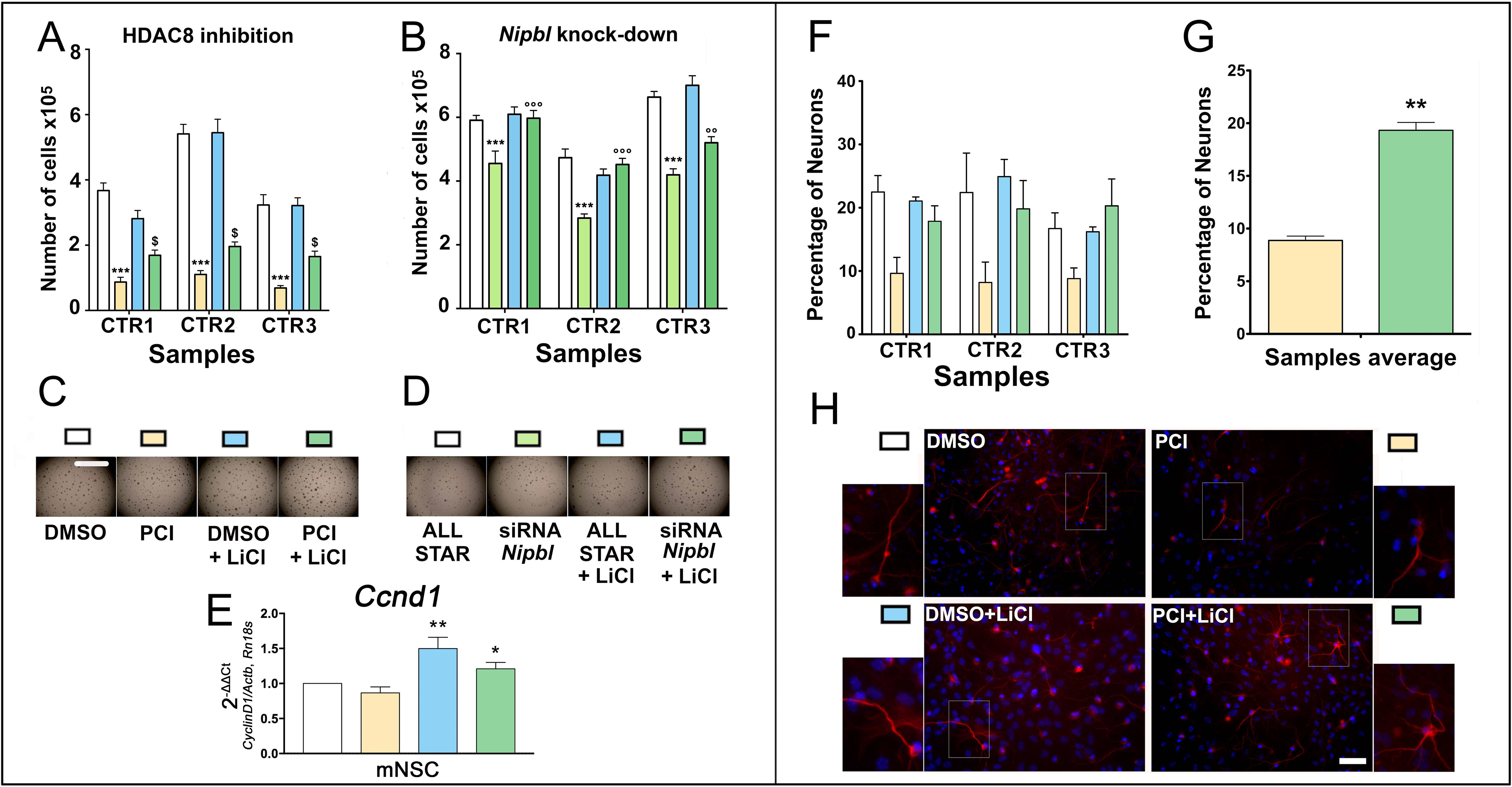
Lithium rescues proliferation and differentiation capabilities in CdLS mouse NSCs through WNT activation. (A) Analysis of the effects of PCI34051 and lithium exposure on the proliferation capabilities of the NSCs. White: NSCs treated with DMSO; yellow: NSCs treated with PCI34051; light blue: NSCs treated with DMSO + LiCl; dark green: NSCs treated with PCI34051 and LiCl. *** p<0.001; $ p<0.05; * DMSO vs PCI34051; $ PCI34051 vs PCI34051 + LiCl. (B) Analysis of the effects of *Nipbl* knock-down and lithium exposure on the proliferation capabilities of the NSCs. White: NSCs treated with AllStar; light green: NSCs treated with siRNA against *Nipbl*; light blue: NSCs treated with AllStar and LiCl; dark green: NSCs treated with siRNA against *Nipbl* and LiCl. *** and °°° p<0.001; * AllStar vs siRNA against *Nipbl*; ° siRNA against *Nipbl* vs siRNA against *Nipbl* + LiCl. (C) Representative pictures of the neurospheres growing in the well during various treatments as in panel A. Scale bar: 2 mm. (D) Representative pictures of the neurospheres growing in the well during various treatments as in panel B. (E) Analysis of the effects of PCI34051 and lithium exposure on the gene expression of *Ccnd1* in the NSCs. White: NSCs treated with DMSO; yellow: NSCs treated with PCI34051; light blue: NSCs treated with DMSO + LiCl; dark green: NSCs treated with PCI34051 and LiCl. ** p<0.01; * p<0.05; ** DMSO + LiCl vs PCI34051; * PCI34051 + LiCl vs PCI34051. (F) Analysis of the effects of PCI34051 and lithium on the differentiation capabilities of the NSCs. Cells were immunostained for neuron and nuclei detection. White: NSCs treated with DMSO; yellow: NSCs treated with PCI34051; light blue: NSCs treated with LiCl; dark green: NSCs treated with PCI34051 and LiCl. (G) Comparison of the effects of PCI34051 and PCI34051 + LiCl of NSCs differentiation. ** p<0.01; * PCI34051 vs PCI34051 + LiCl. (H) Representative staining of the differentiated NSCs. Differentiated neurons were labelled with β-tubulin III antibody (red) and nuclei were labelled with DAPI (blue). Scale bar: 50 μm.

Selective chemical HDAC8 inhibition also impairs NSCs differentiation toward the neuronal lineage (13). Hence, we sought to mitigate the adverse effects of chemical WNT activation in this context. Lithium exposure induced a significant increase in differentiated Tuj1-positive cells (neuronal lineage) in CdLS NSCs, which is comparable to the differentiation rate observed in controls (Fig. 3F-H), in essence counteracting the inhibitory effect of PCI34051.

### Lithium restores proliferation rate in CdLS lymphoblastoid cell lines

To evaluate possible ameliorative effects of lithium treatment in patients *in vitro*, we studied lymphoblastoid cell lines from CdLS patients and healthy donors. As expected (27), proliferation appeared to be reduced in patient-derived cells, when compared to control lines. Interestingly, upon lithium exposure proliferation of CdLS cells was significantly increased (Fig. 4A and Supp. Fig. 3A-B). In fact, CdLS lines exposed to a range of lithium chloride (LiCl) concentrations (1 mM, 2.5 mM and 5 mM) showed increased proliferation, when compared to untreated CdLS cells, or to treated healthy donors (HD) cells (Fig. A and Supp. Fig. 3A-B). Importantly, these effects on proliferation were associated with changes in cell death. In fact, assay analyses showed a decrease in death cells derived from patients treated with lithium (Fig. 4B-C). Remarkably, the lithium dependent increase in proliferation correlated with a decrease in cell death in patient-derived cells, both compared to untreated cells or to HD control lines. To confirm that such ameliorative effects were mediated by WNT activation, we evaluated *Cyclin D1* expression, a known target of WNT canonical pathway. In CdLS cells, *Cyclin D1* showed a trend of increased expression following lithium exposure in CdLS cell lines (Fig. 4D-E). Finally, the same positive effects were observed exposing the cell lines to other chemical compounds (i.e. BIO, IQ-1, deoxycholic acid, CHIRR99021) with known WNT-activation effects (Supp. Fig. 3C-F).

**Figure 4.**
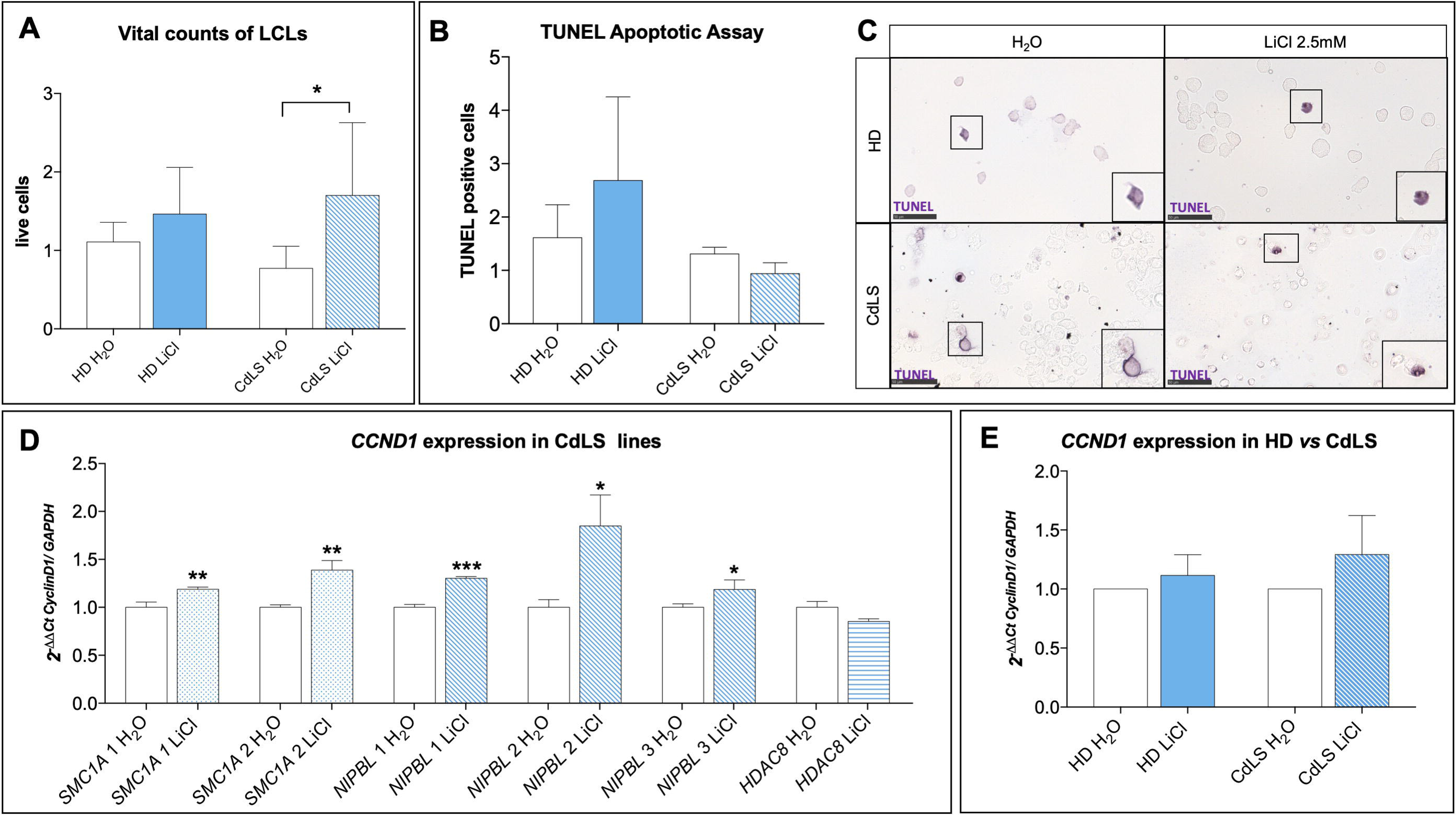
Lithium rescues cell survival in CdLS LCLs through WNT activation. (A) Proliferation of patient-derived cells (CdLS H_2_O, white bar) is reduced compared to the proliferation rate of controls (HD H_2_O, white bar). All CdLS lines (striped bar) exposed to lithium chloride (2.5 mM) showed increased proliferation compared to untreated CdLS cells (CdLS H_2_O, white bar) and compared to treated HD cells (solid blue bar). (B, C) TUNEL assay was used to evaluate cytotoxicity in CdLS LCLs compared to healthy donors (HD). (B) Upon lithium exposure, CdLS cell lines (blue striped bar) show a decrease in cell death compared to untreated CdLS cells (CdLS H_2_O, white bar) and treated HD cells (blue solid bar). On the axis are reported: the experimental groups (x-axis), and numbers of TUNEL positive cells at 24hours of lithium exposure, normalized on water/vehicle (y-axis). Bars express mean ± SEM. (C) Examples of positive TUNEL cells. Images were taken at 40X, while insets display magnification of the white square (80X). Scale bar represents 50 μm. (D-E) *CyclinD1* gene expression was significantly increased upon lithium exposure (2.5 mM) in LCLs, especially in CdLS lines (*SMC1A*: blue dotted bars, *NIPBL*: blue oblique bars) except for *HDAC8* (blue horizontal bar). (D) CdLS lines are shown separately. (E) In pulled CdLS lines (blue striped bar) *CyclinD1* gene expression was increased upon lithium exposure (2.5 mM) compared to controls (blue solid bar). Data are shown as fold change, calculated as 2^−ΔΔCt^ ± SD. p ≤ 0.05 (*), p ≤ 0.01 (**), p ≤ 0.005 (***).

## DISCUSSION

Basic research on cohesins, whose haploinsufficient inactivation causes cohesinopathies such as CdLS, has had a growing impetus in recent years since the role of cohesins on gene expression regulation was identified (28). This function adds to those which have long been extensively studied in the control of DNA damage and in the regulation of chromatid separation during the cell cycle, which have been defined as the “canonical role” (7). Several studies have attempted to clarify which cohesin function is most commonly implicated in CdLS pathogenesis: thanks to *in vitro* and *in vivo* experimental studies, it is now currently believed that the “non-canonical” functions are mainly altered by pathogenic mutations (7, 29).

Control and regulation of different gene pathways could account for the heterogeneity of the clinical involvement of patients with CdLS (30): in fact, these patients have multi-organ malformations, as well as mental retardation and medical complications depending on the degree of disability.

In a previous work of our group (8), aimed at defining the cohesins expression in the different mammalian tissues, a significant expression of cohesins was demonstrated in the developing hindbrain and in the adult cerebellum. The share of replicating cells in the adult cerebellum is limited, so it is possible to assume that the role of cohesins in maintaining the correct adhesion of sister chromatids is marginal in this context, and that there may be involvement of the “non-canonical” role of these proteins. Our previous studies on a *D. rerio* embryonic model of CdLS found a high expression of *nipped-b* in the hindbrain, the structure from which the cerebellum originates (12). Morphological alterations have been identified precisely at the development of the rhombencephalon and a disruption of the WNT pathway was shown (11). It would be extremely significant to pinpoint the molecular cascade regulated by cohesins in differentiating CNS cells, and cohesin contribution to proper hindbrain and hindbrain-derived structures patterning. The difficulty in performing an accurate and objective neurological examination in a CdLS patient, stemming from the seriousness of the cognitive delay, explains the apparent absence of symptoms referable to anomalies affecting the cerebellum. The CNS anomalies reported in the scientific literature often concern single case reports or cases with a small number of patients. In the context of rare conditions, it is in fact difficult to have large cohorts of patients and, in particular, for problems and complications that do not involve all affected individuals. Previous studies are represented on small cohorts, i.e. 8 and 15 patients (31, 32). The cohort presented in this study, therefore, represents the largest ever described, with 66 patients undergoing a brain MRI examination. The number of neuroradiological tests is limited considering the original cohort (155 patients), but this is in line with what has also been highlighted by recent international guidelines that recommend performing brain MRI only in the case of manifest neurological symptoms (1). The evaluation of the MRI reports showed that in a considerable percentage of patients, morphological alterations of the cerebellum are present. Cerebellar anomalies are, in fact, visible in 24.5% of the patients whose MRI data are available. These primarily include hypoplasia of the cerebellar vermis. The finding of cerebellar hypoplasia in patients with genetic syndromes is relatively frequent and, to date, there is much debate regarding the clinical neurological consequences that this malformation may entail (33). Over the years, pathological findings of the cerebellum have been frequently linked and held responsible for the development of behavioral problems attributable to the autistic spectrum: many autopsies of patients with ASD have demonstrated a decrease in the number of Purkinje cells and hypoplasia of the cerebellar vermis or cerebellar lobes (34). The analysis of our cohort shows a statistically significant correlation between CNS anomalies, in particular the structures of the posterior cranial fossa of rhombencephalic derivation, and the presence of autistic features. Analyzing the case history of 15 patients, the Kline group (32) had not reported a correlation between behavioral problems and CNS anomalies, perhaps precisely because of the small number of patients evaluated. Our finding represents additional support of the potential relationship between cerebellar anomalies and autism.

We then used *Drosophila* to model CdLS brain abnormalities *in vivo*. We have shown how the activation of WNT through the administration of lithium helps recover the morphology and structure of brain mushroom bodies also in adults. Our data show how molecular recovery can lead to macroscopic structural improvement. It would be interesting to assess possible feedback in circadian behavior, as it has also been associated with WNT and mushroom bodies patterning in flies (35) and it is known to be disturbed in CdLS patients (36). It is known that WNT has a fundamental role in the development of the CNS, specifically the development of the rhombencephalon (4). Hence, we assessed lithium effects in CdLS mouse NSCs (13). Upon *Nipbl* knockdown by siRNA or Hdac8 inhibition by PCI34051 administration, we observed decreased proliferation and increased apoptosis. By administering lithium, proliferation/death rate was restored to control levels. Furthermore, lithium exposure restores neuronal differentiation capabilities. We also confirmed that this effect was mediated by WNT activation, through the observed increase of *Ccnd1* expression. Finally, we studied LCLs derived from CdLS patients (with pathogenic variants in several causative genes) which were compared to healthy controls. In this experimental model, we confirmed results on CdLS-WNT pathway relationship.

The correlation between rhombencephalic anomalies and autistic features suggests that the use of lithium may represent a therapeutic strategy aimed at improving the behavioral disability of CdLS patients. As previously described, the WNT pathway modulated by lithium, has an essential role in the development of the CNS and a reactivation of the signaling of this pathway determines effects in terms of proliferation and differentiation that could potentially improve cellular deficits caused by an altered timing and differentiation success. We believe that it would be fundamental to study at cellular level in the CNS possible defects and their correlation with neurological signs. A clinical trial has proposed lithium therapy for the treatment of fragile X syndrome showing promising results (37) compared to the effects on behavioral problems. Another work showed the efficacy of lithium as a mood stabilizer in two patients with *SHANK3* mutation who had been diagnosed with an ASD (38). Furthermore, hypoplasia of the cerebellum has been linked to psychomotor retardation of varying degrees even in patients without autistic symptoms (39). In our case history, we find a statistically significant correlation between CNS abnormalities and severe mental retardation but not a relationship with specific abnormalities. Our group (40) has recently developed a prognostic score capable of predicting the severity of intellectual impairment of patients on the basis of clinical features detectable in the first 6 months of life. Should the data emerging from this work be confirmed, the presence of CNS anomalies could represent one of those markers, indicative of a more unfavorable prognosis in terms of intellectual impairment.

In conclusion, in the present study we have further analyzed the significant role of the canonical WNT pathway in the development of the CdLS adverse phenotype. We have demonstrated with *in vivo* and *in vitro* models how lithium, a known WNT-activator, ameliorates phenotypes associated to CdLS by modulating WNT pathway. The comparison with the clinical data revealed a greater prevalence of anomalies of the central nervous system of rhombencephalic derivation, in full consistency with the data of the experimental models. Our findings should be confirmed, possibly through an *ad hoc* prospective study, designed with predefined neuro-behavioral parameters functionally assessed and a homogeneous analysis of the MRI images. This could allow for assessing the use of prognostic brain MRI data with respect to prevalence of behavioral disorders and severity of intellectual impairment.

## MATERIALS AND METHODS

### CdLS imaging

#### MRI brain analyses

The cohort of patients was composed of 155 patients (10 Polish and 145 Italian) with a diagnosis of CdLS according to diagnostic criteria (41). 51.6% were males, and age spanned between 1 month and 53 years (mean 19.83 years old). Exclusion criteria were genetic rearrangement not involving known causative genes (Supp. Fig. 1B). Behavioral and cognitive parameters were assessed by specialized clinicians. Among them 66 patients had performed brain MRI (Fig. 1A).

### Drosophila melanogaster

#### Husbandry

Flies were reared on cornmeal, yeast and molasses media in agar at 25°C. For all experiments, heterozygous *Nipped-B^407^* haploinsufficient mutants (*yw; Nipped-B^407^ . pw^+^ / CyO, Kr-GPF*) (23) and *yw* controls were grown at the same time with the same batch of food preparation. The loss of function allele *Nipped-B^407^* was obtained by γ-ray mutagenesis and previously described (42).

#### Animal treatments

LiCl was added during the cooling step (temperature below 60°C), of the food cooking process, to reach a final concentration of 100 mM; this concentration was determined according to the literature (35, 43–45). Treatments’ outcomes have been assessed after 15-20 days of culturing on the offspring of the fly population exposed to control or supplemented food.

#### Mushroom bodies immunostaining

The protocol for dissection and staining has been optimized from the literature (46). *Drosophila* brains were dissected in 1% Triton-X 100 in phosphate buffered saline (PBT), then fixed in paraformaldehyde (PFA) 4% for 1 hour and stained with mouse anti-FasII 1:20 (clone 1D4; Developmental Studies Hybridoma Bank) for 2 nights, followed by incubation with Alexa-488 anti-mouse secondary antibody (1:1000 - ThermoFisher Scientific, #A21202) for 1 hour. Between each step, samples were washed with PBT on mild agitation. Brain samples were then whole mounted in glycerol based mounting media and slides were sealed. Mushroom bodies were visualized using a fluorescence microscope (Leica DMRB) and acquired using a digital camera (Leica DFC480).

#### qPCR analysis (*Drosophila melanogaster*)

Total RNA was extracted from 15 flies per genotype using the RNeasy Mini Kit (#74106, Qiagen) following manufacturer’s protocol. RNA was quantified using a NanoDrop™ One (#ND-ONE-W, ThermoFisher Scientific) and 2 μg per sample were used for cDNA synthesis by performing a retro-transcription reaction with the SuperScript™ VILO™ cDNA Synthesis Kit (#11754050, Invitrogen) following manufacturer’s protocol. SsoFast™ EvaGreen^®^ Supermix (#1725201, Biorad) with 500 nmol primers were used for qPCR analyses on a CFX96 Touch Real-Time PCR Detection System (#1855196, Biorad). Primers used were home-made designed and tested (Supplementary Table S2). The experiments were performed with three independent biological samples (n = 3) and carried out with technical triplicates. Data are shown as fold change, calculated as 2^−ΔΔCt^ using *RpL32* as the calibrator gene.

### Mouse Neural Stem cells

#### Neural stem cells preparation

Primary cultures and cultivation of the NSCs from adult mice, their differentiation and their immunostaining were carried out as previously described (47, 48) (Supplementary materials and methods). Cells were maintained in the proliferation state using an optimized medium (proliferation medium PM) described in a previous work (49) as neurospheres that were mechanically dissociated every 4-6 days and replated. Every dissociation represents a passage; in the present study, we used cells cultivated for less than 15 passages.

#### Proliferation assays on NSCs exposed to PCI34051 and lithium

Cells maintained in the proliferative state were treated with PCI34051 25 μM for inhibiting HDAC8. For WNT pathway activation LiCl was used (50). For each experiment, 4 groups were studied: (1) DMSO 1:1000 (as control), (2) PCI34051 25 μM, (3) DMSO 1:1000 + LiCl 3 mM and (4) PCI34051 25 μM + LiCl 3 mM. Three different NSCs cultures were used for each experiment and the experiments were run in triplicate. Cells were counted by dissociating the neurospheres in the well and counting the single cells in suspension to determine the total number of cells in the well.

#### Differentiation assays on NSCs exposed to PCI34051 and Lithium

The differentiation of NSC was accomplished by plating the dissociated neurospheres. 40,000 cells were seeded into a 48-multiwell plate containing one 10 mm coated (Cultrex, Tema Ricerca, Italy) round glass coverslip without epidermal growth factor (EGF) for 2 days, then a medium containing 1% of fetal calf serum was used.

During this step, the treatment with dimethyl sulfoxide (DMSO) (1:1000 as control), PCI34051 (25 μM) or LiCl (3 mM) and PCI 34051 (25 μM) + LiCl (3 mM) was also performed. Differentiation was reached after 7 days at 37°C, 5% CO_2_ (13, 51). Cells were then washed once with PBS and fixed with 4% paraformaldehyde (PFA) for 10 min at RT, then kept in PBS until the staining. Two-way ANOVA analysis of the effect of different combinations of PCI34051, and LiCl on NSCs differentiation was carried out.

#### Immunostaining of differentiated NSC

To assess the differentiation of treated NSCs, we used an approach previously described (13) based on antibodies against and β-Tubulin III (Tuj1 1:250, Immunological Sciences AB-10288) and Glial Fibrillary Acidic Protein (GFAP 1:250, Immunological Sciences AB-10635). The cells immunolabeled with β-Tubulin III were counted and the percentage of these cells was calculated by dividing for the overall number of cells (counting the nuclei stained with DAPI).

#### Silencing of proliferating NSCs

Silencing of *Nipbl* was performed using the same protocol described for HDAC8 (13). We used Qiagen siRNA (Flexitube siRNA 5 nmol cat. nr. SI00853111 and cat. nr. SI00853118) and AllStars Negative Control siRNA (cat. nr. 1027280). For allowing siRNA to penetrate into the cells, we used a specifically designed Qiagen Transfection Reagent, HiPerFect.

LiCl treatment was used to assess possible ameliorative effects. To this end, 4 experimental groups were established: (1) control with scramble miRNA (Allstar); (2) *Nipbl*-siRNA obtained using a mix of two different siRNAs at the total concentration of 20 nM (13); (3) control with LiCl treatment (AllStar + LiCl); and (4) *Nipbl*-knockdown with LiCl. Three different NSCs cultures were used and the experiments were performed in triplicates. Cells were counted dissociating the neurospheres in the well, counting the single cells suspension and calculating the total number of cells in the well.

#### qPCR (NSCs)

Cells were resuspended in Trizol reagent (Sigma-Aldrich); subsequently, RNA extraction was performed and contaminating DNA was removed with DNase I (#E1010, Zymo Research) following manufacturer’s protocol. First strand cDNAs were synthesized using SensiFAST™ cDNA Synthesis Kit (#BIO-65054, Bioline) following manufacturer’s protocol. qPCRs were carried out using *TB Green* Premix Ex Taq (Tli RNase H Plus) (#RR420A, Takara Bio Inc., Kusatsu, Japan) and the *Applied Biosystems* StepOnePlus™ *Real-Time* PCR System (Thermo Fisher Scientific, Waltham, MA, United States). Each sample was assayed in technical triplicates and *Ccnd1* levels are expressed as 2^−ΔΔCt^ ± SEM and normalized on values derived from geometric mean of two tested housekeeping genes (*Actb* and *Rn18s*). Both biological and technical triplicates were used and paired groups. The primers used are reported in (Supplementary Table S1). Experiments were performed using RNA from two NSCs colonies cultured in biological triplicates.

### Lymphoblastoid immortalized cell lines

#### Cell culturing

Lymphoblastoid lines (LCL) from patients with different mutations were used: 6 lines from CdLS patients carrying mutations in *NIPBL* (3), *HDAC8* (1) or *SMC1A* (2) (52) (Supplementary Table S3) and 4 lines from healthy donors (HD) serving as controls. Cells were cultured in suspension in RPMI-1640 medium supplemented with 20% fetal bovine serum (FBS), 1% penicillin/streptomycin and were maintained at 37°C in a humidified incubator with 5% CO_2_.

#### Lithium exposure and additional compounds

Cells were exposed to three different concentrations of LiCl: 1 mM, 2.5 mM, and 5 mM or water (vehicle) as control (53, 54). In addition, we used 4 other WNT activators compounds: BIO (6-bromoindirubin-3’-oxime) (0.1 μM, 0.5 μM, and 1 μM); IQ-1 (5 μM, 10 μM, and 20 μM); Deoxycholic acid (5 μM, 100 μM, and 250 μM); and CHIRR99021 (1 μM, 5 μM, and 10 μM) (55–57) (Supp. Fig. 3C-F). Proliferation/death rates were measured counting cells upon trypan blue staining to evaluate the compounds’ effects on viability and cytotoxicity.

#### TUNEL assay

Apoptosis rate in LCLs was evaluated using terminal deoxynucleotidyl transferase (TdT) dUTP Nick-End Labeling (TUNEL) assay, designed to detect apoptotic cells during the late stages of apoptosis, as previously described (13). Briefly, cells were cytospinned on glass slides and then fixed in 4% PFA for 10 minutes at room temperature (RT) and washed three times in phosphate buffer saline (PBS) for 5 minutes. Staining for apoptotic cells was performed using the AP-*In situ* Cell Death Detection Kit (Roche Diagnostics, Penzberg, Germany) following manufacturer’s protocol. Slides were mounted for microscopic imaging with homemade glycerol based mounting media with anti-fading (1,4-diazabicyclo[2.2.2]octane, DABCO). Apoptotic cells were detected with NanoZoomer-XR Digital slide scanner (Hamamatsu, Japan) for counting.

#### Immunofluorescence assay

For assessing cell proliferation rate, Ki67 immunoassay (58) was used. Slides were stained with primary antibody anti-Ki-67 (1:250, #9129 (D3B5) Cell Signaling) overnight, followed by incubation with Alexa-488 anti-rabbit secondary antibody (1:1000, #6441-30 SouthernBiotech) for 2 hours. Slides were mounted for microscopic imaging with EverBrite Hardset Mounting Medium with DAPI (#23004, Biotium) and positive cells were detected and acquired with NanoZoomer-XR Digital slide scanner (Hamamatsu, Japan) for counting.

#### qPCR (Lymphoblastoid cell lines)

RNA was extracted with Trizol (Sigma Aldrich, Italy) following the manufacturer’s protocol. First strand cDNA was synthesized using SensiFAST™ cDNA Synthesis Kit (#BIO-65054, Bioline, Italy) following manufacturer’s protocol. qPCR was carried out using *TB Green* Premix Ex Taq (Tli RNase H Plus) (#RR420A, Takara Bio Inc., Kusatsu, Japan) and the *Applied Biosystems* StepOnePlus™ *Real-Time* PCR System (Thermo Fisher Scientific, Waltham, MA, United States). Primers used are reported in (Supplementary Table S4). Each sample was assayed in technical triplicates and *CCND1* levels were quantified relatively to the expression of *GAPDH* gene. Data are shown as fold change, calculated as 2^−ΔΔCt^.

### Data analysis

NSCs proliferation statistical analysis was performed using student’s t-test and two-way analysis of variance followed by Bonferroni’s Multiple Comparison Test. NSCs immunostaining statistical analysis was performed using the One-way analysis of variance followed by Bonferroni’s Multiple Comparison Test. qPCR data for NSCs were analyzed multiple comparison in one-way ANOVA. qPCR data for *Drosophila* and LCL were analyzed with student’s unpaired t-test. LCL counting were made picking three randomly selected fields per experimental group (20X magnification), and positive cells were calculated by counting within the three fields, by two operators blinded to experimental groups. MRI data from CdLS patients were analyzed using Fisher test. For all the analyses, p ≤ 0.05 (*) was set as statistically significant, p ≤ 0.01 (**), p ≤ 0.005 (***). Graphs were made using Graphpad Prism 7 and figures were assembled using either GIMP-2.1 or Adobe Photoshop CC.

## ACKNOWLEDGMENTS

The authors are grateful to the following fundings: Fondazione Cariplo (2015-0783 to V.M.); Università degli Studi di Milano Intramural Fundings (to V.M. and G.C..); Molecular and Translational Medicine PhD-Università degli Studi di Milano scholarship (to P.G.); Translational Medicine PhD-Università degli Studi di Milano scholarship (to C.P. and E.D.F.); AIRC (Associazione Italiana Ricerca contro il Cancro) Investigator grant 20661 and WCR (Worldwide Cancer Research) grant 18-0399 (to T.V.); Nickel & Co S.p.A and CRC Aldo Ravelli (to VM). Microscopy observations were carried out at The Advanced Microscopy Facility Platform - UniTECH NoLimits - University of Milan. The authors are grateful to Prof. Richard H. Finnell, Dr Jon Wilson and Ms Dawn Savery for helpful comments.

The authors would also like to express their deepest gratitude to CdLS patients and families for constant support and inspiration. The authors thank Susanna Brusa for graphical support.

## AUTHOR CONTRIBUTIONS

V.M. conceived the project and analyzed data; P.G., C.P., D.B., E.D.F. and A.Z. performed the experiments; M.M., J.W., M.I., P.F.A., L.A. and A.S. analyzed patients data; R.A. assisted in confocal microscope imaging; T.V. provided flies reagents and animals; P.G., C.P., M.M., D.B. and V.M. wrote the manuscript; A.C., R.T., T.V. and C.G. provided guidance in the manuscript revision and data interpretation. All authors: approved the manuscript.

## COMPETING INTERESTS

The authors declare no competing interests.

### Compliance with ethical standards

The study is in accordance with the 1964 Helsinki declaration and its later amendments or comparable ethical standards. Written informed consent of patients or caregivers were collected for biological samples studies.

