## Supplementary Information for "Lithium ameliorates Cornelia de Lange syndrome associated phenotypes in experimental models"

**SUPPORTING INFORMATION**

### Supplementary materials and methods

*Neural stem cells preparation*

Two-month-old C57BL/6 male mice were anesthetized by intraperitoneal injection of chloral hydrate [44] and killed by decapitation. The subventricular zone (SVZ), was removed and transferred to a phosphate buffer solution containing antibiotics and glucose (0.6%) at 4 ºC until the end of all the dissections. The tissue was dissociated in an Earl's Balanced Salt Solution (EBSS) (Sigma-Aldrich, St. Louis, MO) containing 1 mg/ml papain (27 U/mg; Worthington DBA, Lakewood, NJ), 0.2 mg/ml cysteine (Sigma-Aldrich, St. Louis, MO) and 0.2 mg/ml EDTA (Sigma-Aldrich, St. Louis, MO), and incubated for 45 min at 37°C. Tissues were then centrifuged at 120 gs, the supernatant discarded and the pellet re-suspended in 1 ml of EBSS and mechanically dissociated using an aerosol resistant tip for 1000 µm. Disaggregated tissue was re-suspended in 10 ml EBSS and centrifuged at 120 gs for 10 minutes. The supernatant was discarded again, and the pellet re-suspended in 200 µL of EBSS. The pellet was dissociated mechanically using an aerosol resistant tip for 200 µm and the cells were resuspended in 10 ml of EBSS and centrifuged at 120 gs for 10 minutes. The supernatant was discarded and the pellet resuspended in 200 µl of DMEM– F-12 based medium [13] (proliferation medium, PM) containing human recombinant Fibroblast Growth Factor (FGF2) and EGF respectively 10 ng/ml and 20 ng/ml (Peprotech, Rocky Hill, NJ, or Upstate Biotechnology, Lake Placid, NY). The cells were plated at 3500 cells/cm$^2$, after 5-7 days spheres were harvested, collected by centrifugation (10 min at 120 *gs*), mechanically dissociated to a single-cell suspension, and re-plated PM. Stem cells used in these experiments were between the 5$^{th}$ to the 15$^{th}$ passage in culture.

*Differentiation assays staining*

Fixed cells were permeabilized with 0.1 % Triton-X in PBS 1X for 10 min at room temperature, then the primary antibodies were added overnight at 4 °C in PBS with 10% normal goat serum (NGS). Secondary antibodies conjugated with fluorophores were used: Alexa-fluor 555 (Goat anti rabbit Immunological Sciences IS20012) and Alexa-fluor 488 (Goat anti mouse Immunological Sciences IS20010). Nuclei were stained with 4′,6-Diamidine-2′-phenylindole dihydrochloride (DAPI) 300 nM [55]. Then the round coverslips were taken from the well, washed in water to remove residual salts and placed (with the surface containing the cells on a coverslip where a drop of 5 µl of DABCO [41] were put.

Images were acquired using an Inverted Microscope ZEISS Axio Vert.A1 and a camera Retiga R3™. The number of positive cells was counted and compared within treatments. Experiments were repeated three times using three different NSC cultures.

*Silencing Nipbl*

One day before the experiment a 48 MW plate was coated with laminin (Synthetic Laminin Peptide, Sigma SCR127) (150 ug/ml) [13].

In order to perform the knockdown of *Nipbl* expression, we used siRNA and transfection reagents from Qiagen. Two validated siRNAs that were able to anneal different regions of the *Nipbl* transcript were selected (Flexitube siRNA 5 nmol cat nr SI00853111 and cat nr SI00853118), as negative control we used AllStars Negative Control siRNA (cat nr 1027280) which has no homology to any known mammalian genes and that has minimal nonspecific effects. To allow the entrance of the siRNA into the cells, we used HiPerFect Transfection Reagent which is specifically designed by Qiagen for the aforementioned siRNAs.

These experiments were carried out using 3 different cultures of NSCs, the experiments were performed in triplicate. Inasmuch, as the entrance of siRNA in neurospheres was demonstrated to be hard, we first plate the dissociated cells (10,000 cells per well) on laminin-coated wells as a monolayer in 500 µl of proliferation medium (PM) for one day. The day after we prepare 100 µl incubation medium made by PM and siRNA (at the chosen concentration) and 1.5 µl HiPerFect Transfection Reagent. This medium was mixed by vortexing (10 sec) and maintained 5–10 min at room temperature (15–25°C) to permit transfection complexes formation. The incubation medium was added dropwise onto the cells swirling the plate gently. The cells were incubated for 3 hours with the transfection complexes at 37 °C 5% $CO_2$ and then 400 µl of PM were added. Twenty hours after the medium was changed with fresh PM and the cells were left grown for 2-3 days. The cells were then harvested, dissociated and counted. We first performed a setup of the experiment assessing the appropriate concentration of the siRNA. Considering previous work [13] we used 20 nM (total siRNA) that was able to reduce the proliferation at higher levels.

*Lymphoblastoid immortalized cell lines*

The cells were cultured in T-25 flasks and were grown at 37 °C, 5% $CO_2$, mechanically dissociated to a single-cell suspension, and re-plated using RPMI medium (20% foetal bovine serum, FBS; 1% streptomycin). Cells were treated for 24 hours during exponential phase of growth and counted at 2 different timepoints: before and after chemical exposure (O, 24 hours).

**Supplementary figures and figure legends**

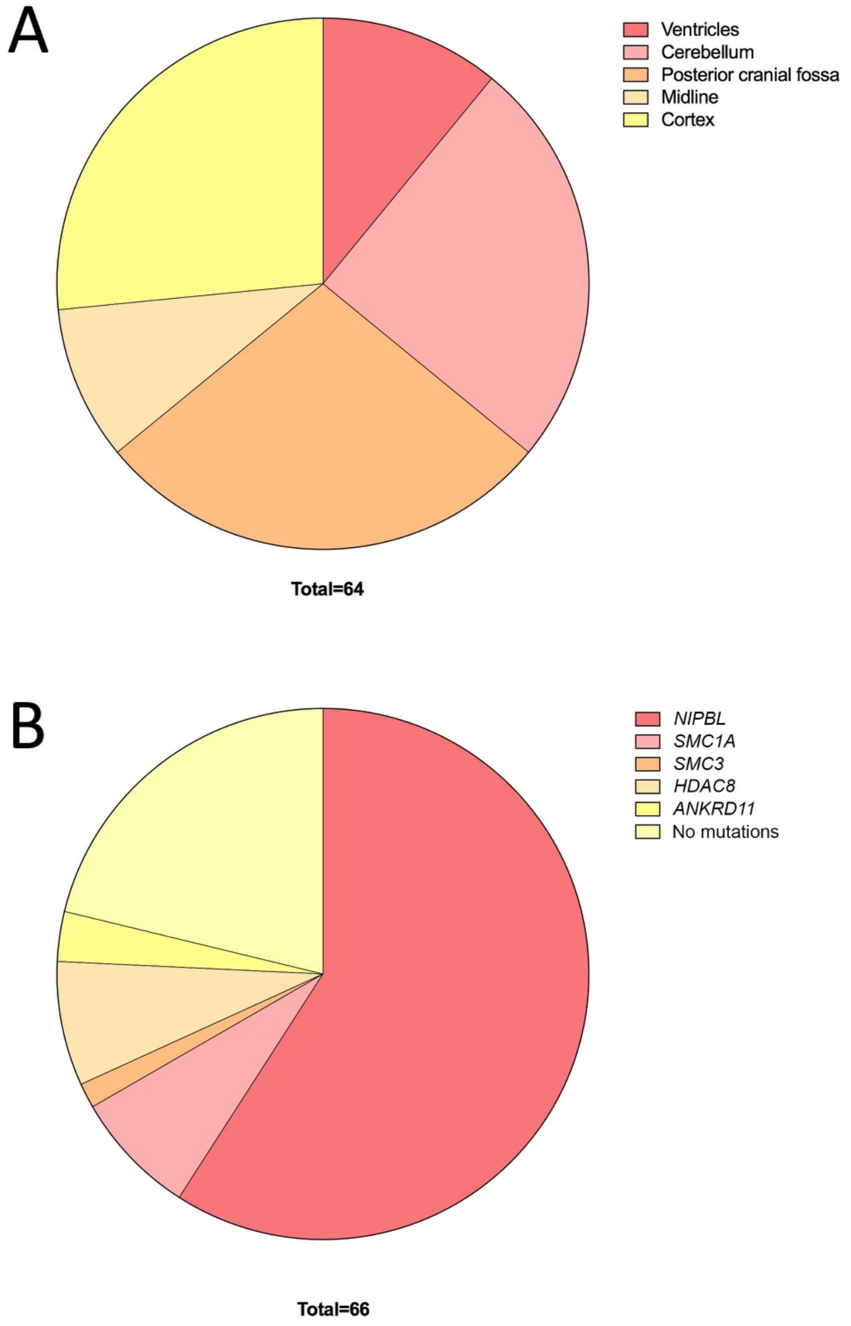


**Supplementary Figure 1. Brain MRI and CdLS patients**

(A) MRI data distribution of tissue anomalies in CdLS patients. (B) Distribution of genetic variants in the CdLS patient cohort.

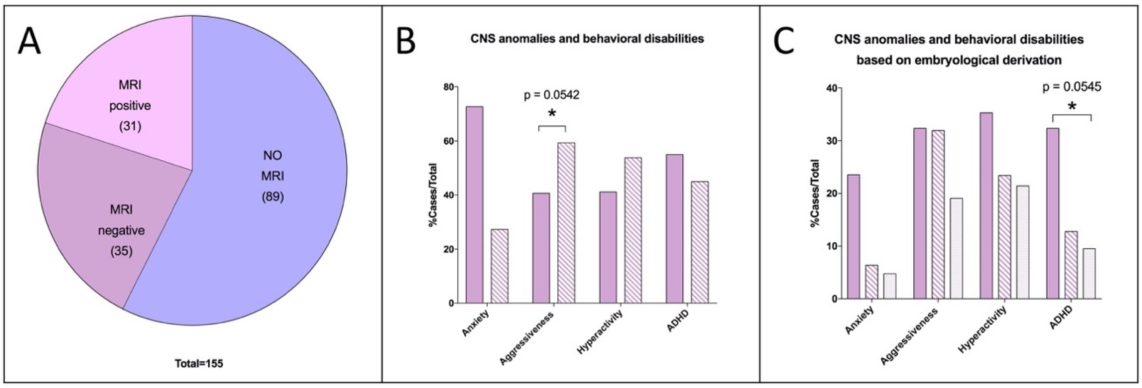


**Supplementary Figure 2. CdLS Brain abnormalities in hindbrain-derived structures correlates with cognitive and behavioral alterations**

(A) MRI data distribution in the cohort of CdLS patients. Purple: no MRI (89/155); mauve: MRI negative (35/155); pink: MRI positive (31/155). (B) CNS anomalies and behavioral disabilities. Solid mauve: MRI negative; striped mauve: MRI anomalies. (C) CNS anomalies and behavioral disabilities based on the embryological derivation. Solid mauve: MRI negative; striped mauve: Rhombencephalon derivation; dotted mauve: Prosencephalon derivation. * $p<0.05$; * vs MRI negative.

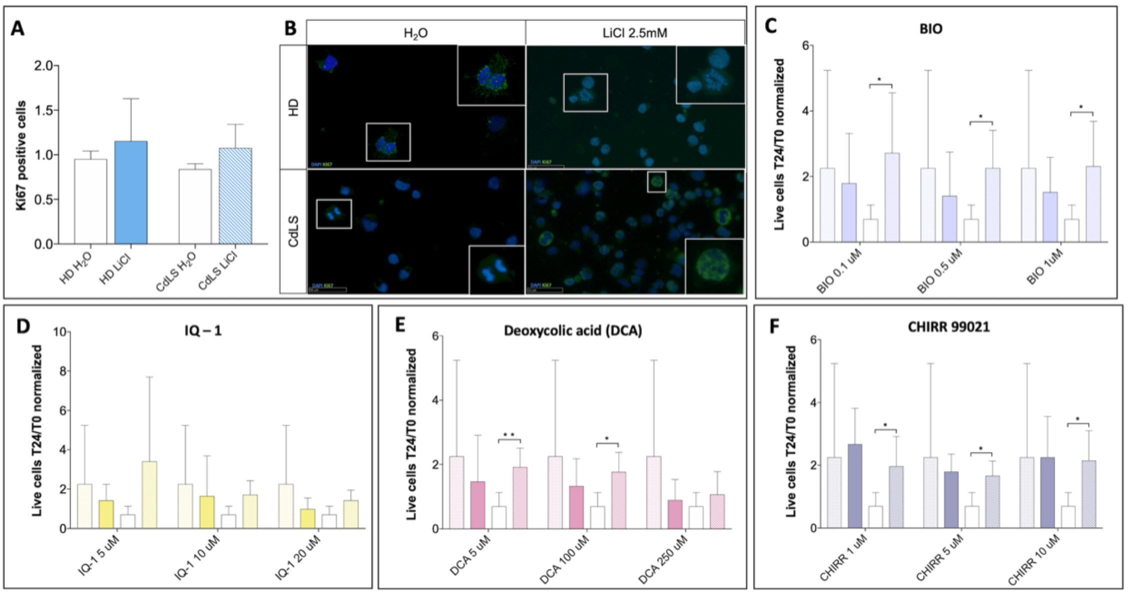


**Supplementary Figure 3. Vital counts of lymphoblastoid cell lines**

(A, B) Proliferation assay using Ki-67 was performed to evaluate the viability in CdLS LCLs compared to healthy donors (HD). (A) In standard conditions, the proliferation of patient-derived cells (CdLS $H_2O$, white bar) is reduced compared to the proliferation rate of healthy donors (HD $H_2O$, white bar). CdLS cell lines (blue striped bar) exposed to LiCl (2.5mM) have an increased proliferation compared with untreated CdLS cells (CdLS $H_2O$, white bar) and treated HD cells (blue solid bar). On the axis are reported: the experimental groups (x-axis), and numbers of cells in proliferation at 24 hours of lithium exposure, normalized on water/vehicle (y-axis). Bars express mean ± SEM. (B) Examples of cells in proliferation following Ki-67 immunoassay. Images were taken at 40X, while insets display magnification of the white square (80X). (C-F) Treatments with others WNT pathway activators are shown (HD untreated: dotted bars; treated HD: solid colored bars). Untreated CdLS cells (white bars) are exposed to different concentrations of the compounds (oblique striped bars). Treatments with (C) BIO 0.1 µM, 0.5 µM, 1 µM; (D) IQ-1 5 µM, 10 µM, 20 µM; (E) Deoxycolic acid 5 µM, 100 µM, 250 µM; (F) CHIRR99021 1 µM, 5 µM, 10 µM are represented. On the axis are reported: the experimental groups (x-axis), and numbers of number of live cells at 24 hours of exposure divided by number of live cells at T0, normalized on water/vehicle (y-axis). * $p< 0.05$, ** $p<0.01$.

| Mouse gene | Primer sequence |
|---|---|
| *Cyclin D1 (Ccnd1)* | Forward: CGTGGCCTCTAAGATGAAGGA |
| | Reverse: CCTCGGGCCGGATAGAGTAG |
| *Actin, beta (Actb)* | Forward: TCCATCATGAAGTGTGACGT |
| | Reverse: GAGCAATGATCTTGATCTTCAT |
| *18S ribosomal RNA (Rn18s)* | Forward: TTGACGGAAGGGCACCACCAG |
| | Reverse: GCACCACCACCCACGGAATCG |

**Supplementary Table S1 - Primers' sequences for qPCR analysis (mouse NSCs)**

| ***Drosophila melanogaster* gene** | **Primer sequence** |
|---|---|
| *Armadillo (arm)* | Forward: TCTGCTGCAACGAAACAACG |
| | Reverse: CTGCATCCGAAAGATTGCGG |
| *Engrailed (en)* | Forward: TATCGCCGCACTTCAAAAGC |
| | Reverse: TTTACAGAGCGGTTGCAAGC |
| *Ribosomal protein L32 (RpL32)* | Forward: ACAGGCCCAAGATCGTGAAG |
| | Reverse: CTTGCGCTTCTTGGAGGAGA |

**Supplementary Table S2 - Primers' sequences for qPCR analysis (*Drosophila melanogaster*)**

| CdLS LCLs | Gene | Exon | cDNA change | Protein change | Mutation type | Clinical features | Reference |
|---|---|---|---|---|---|---|---|
| Sp11 | ***NIPBL*** | 19 | c.4253G>A | p.(G1418E) | Missense | Mild | This work |
| Sp12 | ***NIPBL*** | 4 | c.231-2_231-1delAG | p.(E78Vfs*4) | | Moderate | This work |
| Sp47 | ***NIPBL*** | - | t(5;15);inv(5p) | - | | Severe | This work |
| Sp35 | ***SMC1A*** | 2 | c.173del15 | p.(V58_R62del) | In frame deletion | Moderate | Gervasini et al. 2013 (pt #2) |
| 202 | ***SMC1A*** | 15 | c.2351T>C | p.(I784T) | Missense | Moderate | Gervasini et al. 2013 (pt #5) |
| Sp59 | ***HDAC8*** | 10 | c.1022delG | p.(G341Vfs*33) | Frameshift | Mild | This work |

**Supplementary Table S3 - Lymphoblastoid immortalized lines from patients, clinical features and related mutations**

| Human gene | Primer sequence |
|---|---|
| *Cyclin D1 (CCND1)* | Forward: CTGGAGGTCTGCGAGGAA |
| | Reverse: GGGGATGGTCTCCTTCATCT |
| *Glyceraldehyde 3-phosphate dehydrogenase (GAPDH)* | Forward: GAGTCAACGGATTTGGTCGT |
| | Reverse: TTGATTTTGGAGGGATCTCG |

**Supplementary Table S4 - Primers' sequences for qPCR analysis (Lymphoblastoid cell lines)**
